# Pioglitazone improves deficits of *Fmr1*-KO mouse model of Fragile X syndrome by interfering with excessive diacylglycerol signaling

**DOI:** 10.1101/2020.09.22.301762

**Authors:** Andréa Geoffroy, Karima Habbas, Boglarka Zambo, Laetitia Schramm, Arnaud Duchon, Eric Flatter, Laetitia Fouillen, Julie Zumsteg, Dimitri Heintz, Jean-Louis Mandel, Yann Hérault, Hervé Moine

## Abstract

Fragile X syndrome (FXS), the leading cause of familial intellectual disability, is an uncured disease caused by the absence or loss of function of the FMRP protein. FMRP is an RNA binding protein that controls the translation of specific proteins in neurons. A main target of FMRP in neurons is diacylglycerol kinase kappa (DGKk) and the loss of FMRP leads to a loss of DGK activity causing a diacylglycerol excess in the brain. Excessive diacylglycerol signaling could be a significant contributor to the pathomechanism of FXS. Here we tested the contribution of DAG-signaling in *Fmr1*-KO mouse model of FXS and we show that pioglitazone, a widely prescribed drug for type 2 diabetes, has ability to correct excessive DAG signaling in the brain and rescue behavioral alterations of the *Fmr1*-KO mouse. This study highlights the role of lipid signaling homeostasis in FXS and provides arguments to support the testing of pioglitazone for treatment of FXS.

## Introduction

Fragile X syndrome (FXS) is the leading cause of familial intellectual disability (affecting approximately 1 in 5000 males and 1 in 8000 females) and is associated with cognitive and behavioral deficits that can include anxiety, hyperactivity, hypersensitivity, stereotypies, memory deficits, sleeping problems (1). FXS is due to the transcriptional silencing of the *FMR1* gene encoding the fragile X mental retardation protein (FMRP)(2) or, more rarely, to mutations in FMRP itself (3-5). FMRP is an RNA binding protein associated with numerous mRNA species in neurons (6-10) and in other cell types (11). FMRP has been proposed to control the translation of a number of its mRNA targets in neurons and the alteration of neuronal physiology associated with FXS is linked to an excessive protein translation in neurons resulting from the loss of FMRP control (12). A protein synthesis-dependent synaptic plasticity, group 1 metabotropic glutamate receptor long term depression (mGluR1-LTD), is altered in various areas of the brain (13-15) of *Fmr1*-KO mouse. Accordingly, several downstream effectors of the mGluR1-dependent signaling have been shown to be overactivated (e.g. PKC (16), Ras/MEK/ERK/Mnk (17, 18), PIK3K/AKT (19-21), CB1R (22-24), while others are defective (Rac/PAK (25)). In agreement with these data, the phosphorylation of eIF4E by Mnk (26) and of S6 by ERK (18) is increased and an excess of protein synthesis is observed in vitro and in vivo notably in hippocampal and cortical neurons of *Fmr1*-KO mouse (27, 28). How exactly these alterations result from the loss of FMRP remains unclear. We previously identified that diacylglycerol kinase kappa mRNA (DGKk) is the mRNA species most bound by FMRP in dissociated cortical neurons from new born mice (8). In addition, DGKk expression is severely reduced in brain of *Fmr1*-KO mouse and the lack of FMRP leads to a decreased DGK activity in the neurons of *Fmr1*-KO mouse and in the brain of FXS patients causing a perturbation of diacylglycerol/phosphatidic acid homeostasis (8). Moreover, the knockdown of DGKk in the brain of wild-type mouse recapitulates main *Fmr1*-KO behaviors, and DGKk reexpression in *Fmr1*-KO hippocampal slices rescues the abnormal dendritic spines morphology (8). These data support a model where DGKk dysregulation plays a pivotal role in FXS pathomechanism (29).

A prediction of this model of FXS states that interfering with DGKk dysregulation could be beneficial for FXS condition. Pioglitazone (PGZ), a thiazolidinedione drug approved for type 2 diabetes treatment was shown to activate DGK activity and inhibit the DAG-PKC signaling (30). Thus, PGZ could have a potential to correct some of the defects associated with DGKk loss of activity. In this study, we tested the impact of PGZ in neurons in vitro and on the *Fmr1*-KO mouse to rescue the DAG-PKC signaling-related defects of FXS. We first confirmed that DAG-PKC signaling is excessive in *Fmr1*-KO neurons and PGZ normalized these alterations. Then, after defining that PGZ is efficiently delivered to brain by intraperitoneal (IP) administration route, we showed that PGZ treatment improved several of *Fmr1*-KO main behavioral defects, emphasizing the importance of DGKk dysregulation in FXS and highlighting a possible new intervention mean.

## Materials and methods

### Ethics statement

Animal work involved in this study was conducted according to relevant national comité national de réflexion éthique en expérimentation animale with approval APAFIS#5874-20l6062915583967 v2 and international guidelines (86/609/CEE).

### Animal housing

In vivo experiments were conducted in *Fmr1*-KO2 male mice (E.J. Mientjes et al., 2006). C57BL/6J *Fmr1* +/y males (Janvier Labs, France) were crossed with C57BL/6J *Fmr1* +/- females. At weaning age (4 weeks), male mice were grouped by 3 or 4 individuals from same age and genotype in individually ventilated cages (GM500, Tecniplast, UK), with poplar shaving bedding (Lignocell Select, JRS, Germany), and maintained under standard conditions, on a 12-h light/dark cycle (7h/19h), with standard diet food (standard diet D04, Scientific Animal Food and Engineering, France) and water available ad libitum. Mice in the same cage received the same treatment and were transferred to the phenotyping area the following week.

### Animal treatment

Pioglitazone (E6910, Sigma-Aldrich) was administered at 20 mg/kg/day using several administration routes. Food pellets containing vehicle or PGZ (130mg PGZ/KG D04, Safe diet, Augy, France, based on 4-5 g/day pellet consumption) were given from weaning for three weeks before starting the behavioral experiments and continued during the tests. Subcutaneous implants for 60 days treatment (Innovative Research of America, Sarasota, FL, Cat. No. SX-999) prepared with 30 mg PGZ per pellet or vehicle (based on 60 x 0,5mg/day delivery) were placed under dorsal neck skin by sterile surgery. Intraperitoneal injections of PGZ solution at 3 mg/ml in NaCl 0.9%, DMSO 30% (v/v) (or vehicle) were performed daily starting at 5-week of age for one week prior to starting the behavioral tests and continued during the tests (with injection at end of each day test). Well-being of animals was controlled by daily visual control and weekly weighing.

### Cortical neuron cultures

Cortical neuron cultures were prepared as in (8). Briefly, cortices of WT or *Fmr1*-KO (*Fmr1*^-/y^) E17.5 embryos were dissected in PBS 1X, 2.56 mg/mL D-glucose, 3 mg/mL BSA and 1.16 mM MgSO4, incubated for 20 min with 0.25 mg/mL trypsin and 0.08 mg/mL DNase I and mechanically dissociated after supplementation of medium with 0.5 mg/mL trypsin soybean inhibitor, 0.08 mg of DNase I and 1.5 mM MgSO4. Neurons were seeded on poly-L-lysine hydrobromide-coated six-well culture plates in Neurobasal Medium (NBM, GIBCO) supplemented with B27, penicillin/streptomycin and 0.5 μM L-glutamine. Primary cortical neurons were used for experiments after 8 days of culture at 37°C, 5 % CO2. Where indicated, cultures were treated with addition of pioglitazone solution and/or puromycin at the indicated concentrations and times. After treatment, cells were immediately washed with ice-cold PBS and lysed in 4X Laemmli buffer.

### Pioglitazone measurement in brain

Mouse brains (200 mg frozen ground homogenate) were homogenized mechanically with 1:1 w/v H_2_O and then diluted at 1/3 v/v in H2O. Rosiglitazone (Sigma, 2.5ng solubilized in 5μL DMSO/H2O 1:1, v/v) was added as internal standard and 500μl methanol was added to 95μl of brain homogenate. Samples were vortexed for 5min at maximum speed and centrifuged for 20min at 14.000 g at 4°C. 300μL of supernatant (equivalent to 15μg tissues) were transferred to new collecting tube. Samples for establishing standard curve of PGZ were prepared in same conditions by adding 1000ng, 500ng, 250ng, 125ng, 62.5ng or 31.25ng PGZ in (Sigma). Rosiglitazone and PGZ were identified and quantified by UPLC–MS/MS. Quantitative profiles were analyzed using an EVOQ Elite LC-TQ (Bruker Daltonics) equipped with an electrospray ionization source and coupled to an HTC Pal-xt (Bruker Daltonics) and an Advance UHPLC system (Bruker Daltonics). Chromatographic separation was achieved using an Acquity UPLC BEH C18 column (100 × 2.1 mm, 1.7 μm; Waters) and pre-column. The mobile phase consisted of (A) water and (B) methanol, both containing 0.1% formic acid. The run started by 2 min of 95 % A, then a linear gradient was applied to reach 99 % B at 10 min, followed by isocratic run during 1,5 min. Return to initial conditions was achieved in 1 min and equilibrate for 2,5 min, with a total run time of 15 min. The column was operated at 40°C with a flow rate of 0.35 ml/min, injecting 5 μL samples. Nitrogen was used as the drying and nebulizing gas. The nebulizer gas flow was set to 35 L/h, and the desolvation gas flow to 30 L/h. The interface temperature was set to 350°C and the source temperature to 300°C. The capillary voltage was set to 3.5 kV; the ionization was in positive mode. Low-mass and high-mass resolution was 2 for the first mass analyzer and 2 for the second. Data acquisition was performed with the MS Workstation 8 software and data analysis with the MS Data Review software. Absolute quantifications were achieved by comparison of sample signals with dose– response curves established with pure compounds. The MS/MS MRM transitions were 357>134.10 and 357> 339 for the pioglitazone and 358>135 and 358>341.2 for the rosiglitazone.

### DAG measurements

Mouse brains (100 mg frozen ground homogenate) were homogenized with 1 ml H_2_0 and extracted by vortexing with 2 ml of chloroform/methanol 2:1 (v/v) and 10 μl of synthetic internal standard (DAG 15:0/15:0, Sigma Aldrich), sonicated for 30 s, vortexed, and centrifuged. The lower organic phase was transferred to a new tube, and the upper aqueous phase was reextracted with 2 ml of chloroform. Organic phases were combined and evaporated to dryness under nitrogen. Lipid extracts were resuspended in 50 μL of eluent A, and a synthetic internal lipid standard (DAG 17:0/17:0, Nu-Check Prep) was added.

LC-MS/MS (MRM mode) analyses were performed with mass spectrometer model QTRAP 5500 (ABSciex) coupled to a LC system (Ultimate 3000; Dionex). Analyses were achieved in positive mode. Nitrogen was used as curtain gas (set to 20), gas1 (set to 25), and gas2 (set to 0). Needle voltage was at +5,500 V without needle heating; the declustering potential was set at +86 V. The collision gas was also nitrogen; collision energy was adjusted to +34 V. Dwell time was set to 3 ms. Reversed-phase separation was carried out at 30 °C on a Phenomenex Luna 3u C8 column (150 mm × 1 mm, 3-μm particle size, 100-Å pore size). Eluent A was ACN/MeOH/H2O (19/19/2, v/v/v) +0.2% formic acid +0.028% NH4OH, and eluent B was isopropanol +0.2% formic acid +0.028% NH4OH. The gradient elution program was 0–5min, 15% (v/v) B; 5–35min, 15–40% B; 35–40 min, 80% B; and 40–55 min, 15% B. The flow rate was 40 μL/min, and 3-μL sample volumes were injected. The relative levels of DAG species were determined by measuring the area under the peak, determined by using MultiQuant software (Version 2.1; ABSciex) and normalizing to the area of the DAG internal standard.

### Immunoblot analyses

Immunoblot analyses were performed on 14 days-old E19 cortical primary neurons lysed in 4X Laemmli buffer (equivalent to 15 μg total proteins) or brain homogenate (30mg of frozen ground total brain) solubilized in RIPA lysis buffer (140 mM NaCl, 0.5% sodium deoxycholate (w/v), 1% Nonidet P-40 (v/v), 0.1% SDS (w/v), and protease inhibitors (Roche Applied Science), lysed by three cycles of freezing/thawing, centrifuged at 4°C for 30 min at 15,000 g. Brain extracts (equivalent to 15 μg total proteins) were mixed with an equal volume of 2X Laemmli buffer. Immunoblotting were performed as described previously (8). Proteins were denatured by heating the mixture for 5 min at 95°C, and then resolved by 10% SDS-PAGE. Separated proteins were transferred using a Mini Trans-Blot (Biorad) cell onto PVDF Immobilon P membrane (Millipore). Membranes were blocked for one hour with TBS-T 1X (Tris-Buffer Saline, pH 7.4 and 0.1% Tween-20 v/v) containing 5% BSA (w/v) or 5% nonfat dry milk. Membranes were incubated overnight at 4°C with primary antibodies diluted in TBS-T buffer containing 5% w/v BSA or milk as follows: anti-FMRP (1C3, 1/10,000, in house generated antibody) anti-p-EIF4e (Ser209, 1/1000 BSA, #9741, Cell Signaling), anti-EIF4 (1/1000 BSA, #9742, Cell Signaling), anti-p-PKC pan (βII Ser660, Rb, 1/1000 BSA #9371, Cell Signaling), anti-PKCα (1/2000 BSA, sc-208, Santa Cruz), anti-puromycin (12D10, 1/2000, Sigma-Aldrich), anti-S6 (5G10, 1/1000 #2217S, Cell Signaling), anti-p-S6 (Ser235/236, 1/1000, 2211S Cell Signaling), anti-GAPDH (MAB374, 1/10000, Merck) was used as an internal standard. Membranes were washed in TBS-T buffer and then incubated for an hour at room temperature with the corresponding horseradish peroxidase-conjugated preadsorbed secondary antibody (1/2000, blocking solution corresponding, Molecular Probes). Membranes were washed in TBS-T buffer. Immunoreactive bands were visualized with the Immobilon Western Chemiluminescent HRP Substrate (Millipore, cat. WBKLS0100). Immunoblot pictures were acquired using LAS600 GE Amersham and density of the resulting bands was quantified using ImageJ and statistical significance assessed using repeated measures analysis of variance (ANOVA) with Fisher’s post hoc comparisons (GraphPad Prism 7 software, La Jolla, CA, USA).

### RT-qPCR

Total RNAs were extracted with standard Trizol protocol (Trizol, Ambion) from 14 days-old E19 cortical primary neurons (equivalent to one well of 6-well plate) or ground brains (30mg). Total RNA (0,5μg) was treated with DNase (TURBO DNA-free™ Kit, Ambion) following manufacturer instructions. First strand cDNA was synthesized with SuperScript II Reverse Transcriptase kit (Invitrogen) following manufacturer instructions used with random primer mix. qPCR was performed on cDNA dilutions of 1/20 in the presence of 7.5-pmol primers with QuantiTect SYBR Green PCR Master Mix (Qiagen) on a Lightcycler 480 (Roche), using specific primers (Table S1). Relative quantifications were performed with Ribosomal Protein Lateral Stalk Subunit P0 (Rplp0) and Actb as internal standards. Results were expressed as arbitrary units by calculating the ratio of crossing points of amplification curves for cDNAs and internal standard by using the LightCycler 480 software (version LCS480 1.5.0.39, Roche Diagnostics). Fold change between groups were determined by using the following 2e−ΔΔct method. Error rates were calculated with square root ((SD control/SD treated)^2^ + (fold change)^2^).

### Behavior analyses

One week after starting of PGZ treatment behavior tests were conducted to evaluate locomotor activity and anxiety, cognition, autistic-like behaviors and sensori-motor gating. Animals were transferred to the experimental room 30 min before each experimental test. The tests were performed in the following order: open field, novel object recognition, rotarod, stereotypy, sociability, actimetry, pre-pulse inhibition, audiogenic seizures (**Fig. S1**). After each test, material was cleaned with water, 50% ethanol, water, and then dried with paper towels to minimize olfactory cues. IP treatment was applied after behavioral sessions to minimize impact of injection with tests. Behaviors were either analyzed automatically or by an observer blind to the conditions.

### Open field

Locomotor activity and anxiety of animals were evaluated in an open field test. Animals were allowed to freely explore the new environment of plexiglass square arena (44 x 44cm x 16 cm, Panlab Harvard apparatus IR ACTIMETER, Bioseb) at a luminosity of 150 +/- 15 lux. Activity was recorded for 30 min using ActiTrack software (version 2.55). Infra-red lasers at bottom and top of arena recorded vertical and horizontal activity. For the analysis, the arena was divided between a 10-cm wide peripheral zone and a central 30cm x 30cm zone to evaluate locomotion, exploration, hyperactivity and anxiety. Distance moved and time spent in center or periphery zones, velocity, speeds and rearing were scored. Parameters were quantified in cumulative (0-30min) and windows (0-10min, 10-20min and 20-30min).

### Novel Object recognition

Learning and memory abilities of mice were evaluated using the novel object recognition test. The test was performed at a luminosity of 70 +/- 5 lux in the plexiglass square arena used for open field test (44 x 44cm x 16 cm, Panlab Harvard apparatus IR ACTIMETER, Bioseb), considered as a habituation session 24h before training session. First part of the test consisted of a learning task. Two identical objects were presented to animals. Marbles and dices used for the test were randomly placed at defined objects positions. Exploratory activity was recorded for 10min. 24 hours later, second part of the test consisted of a memory recall. Two objects were presented to animals at the same random position (left/right) with a new object replacing one of the learning session objects (next/familiar). Exploratory activity was recorded for 10min. For each session, sniffing time spent on objects and numbers of sniffing contacts were manually scored, defined as <2 cm distance between the nose and the object. Memory capacity was defined by calculated discrimination index (percentage of exploration of the novel object) as the proportion of time that animals spent investigating the novel object minus the proportion spent investigating the familiar one in the testing period: Discrimination Index = [(Novel Object Exploration Time – Familiar Object Exploration Time)/ Total Exploration Time] × 100.

### Rotarod

Locomotors coordination was evaluated during 4 successive session-tests performed at a luminosity of 115 +/- 15 lux on a rotarod system composed of a rotating plastic bar covered by grey rubber foam (5 cm in diameter, Panlab Harvard apparatus LE 8200, Bioseb). The animals were first habituated to stay on the rod for 30 s at a constant speed of 4 rotations per minute (rpm), facing to the direction of rotation. The test was performed on the rotating rod with acceleration steps from 4, 10, 16, 22, 28, 34 to 40 rpm over 5 minutes. First session was considered as a training session, followed by 3 trial sessions separated by 10 min resting. The test was stopped when the mouse fell or when there was more than one passive rotation. The latency to fall and the maximal speed before falling were recorded.

### Stereotypy

Stereotypy was evaluated on animals isolated in a new home cage (GM 500 techniplast, floor area 501 cm^2^ at a luminosity of 70 +/- 5 lux) with sawdust bedding for 10min and measuring exploratory and care activities. Number of rearing or digging events and grooming were scored.

### Free sociability

Sociability was performed at 70 +/- 5 lux in home-cage (GM 500 techniplast, floor area 501 cm^2^) for 10min. For the test two animals were isolated in a new home cage with sawdust bedding for 10min. Two social partners were conspecific of the same group (genotype, treatment), housed in same conditions (4-5 individuals per cages) but were non-siblings and housed in different cages. Number of contacts and times spent in contact were scored for affiliative behaviors (sniffing or touch any part of the body, allogrooming, followings or traversing the partner’s body by crawling over/under it) and aggressive behaviors (attacks, offensive or defensive postures, tail rattling).

### Actimetry

Locomotor and circadian activities were measured in an actimetry recording systems coupled to BMB software (Imetronic). Mice were placed in activity boxes (25 x 20 x 15 cm). Infra-red lasers at the bottom and the top of the square inform about vertical and horizontal activity when cut by mouse body (15 mm for horizontal activity and 30 mm for vertical activity). Activity was recorded during 24 h, starting at 06:00 pm (1h before light switch). Rearing events, front- and back-yard activities, round trips, food and water consumption were recorded.

### Pre-Pulse inhibition (PPI)

PPI was measured using acoustic startle test with SR-LAB apparatus and software (San Diego Instruments, USA). Mice were placed individually in the PPI test plexiglass tube (12,7cm long, 3,8cm internal diameter, 5,0cm external diameter) of the startle box (constant luminosity of 70 +/-5 lux and white noise background). Habituation phase was performed for 5 min. Then mice were exposed to pulses of white sound of 20 msec duration and varying intensity: +6, +12, +18, and +24 dB over background levels (namely 72, 78, 84, and 90 dB). Each intensity was applied eight times, in a randomized order with variable intervals (10–20 sec) between the onset of each pulse. A total of 130 readings of the whole-body startle response were taken at 0.5-msec intervals (i.e., spanning across 65 msec), starting at the onset of the pulse stimulus. The average amplitude (in arbitrary units) over the 65 msec was used to determine the stimulus reactivity and further averaged across trials.

### Statistical analyses

Data were analyzed using an analysis of variance (ANOVA) with Fisher’s post hoc comparisons. ANOVA with repeated measures analysis were carried out when required by the experimental plan to assess complementary statistical effects. In some designs, statistical analysis was performed using Student’s t-tests. For all analysis p-value p<0.05 was considered to indicate significance. All analyses were performed using Statview 5 software for Windows (SAS Institute, Berkley, CA, USA) or GraphPad Prism (GraphPad Software, La Jolla, CA, USA) software.

## Results

### DAG signaling is excessive in *Fmr1*-KO neurons and is normalized by DGK-agonist pioglitazone (PGZ)

We previously showed that the level of second messenger diacylglycerol (DAG) is increased in *Fmr1*-KO cortical neurons and in cerebellum of FXS patient, in agreement with the observed loss of DGKk synthesis in FXS condition (8). DGKk is one of the ten DGK enzymes that control DAG homeostasis (31). Among the first direct effectors of DAG are protein kinase C enzymes (PKC), which are activated through sequential phosphorylation steps (31), notably at carboxy-terminal hydrophobic site Ser660 (32). In agreement with an increase of DAG-induced PKC activation, we measured an elevation of PKC phosphorylation in cultured cortical neurons from *Fmr1*-KO mice compared to their wild-type (WT) littermate control cultures (**Fig. 1A**). As a possible mean to interfere with or compensate for the loss of DGKk in *Fmr1*-KO cortical neurons, we tested the impact of DGK-agonist pioglitazone (PGZ), a molecule with PPARg agonist activity reported to inhibits DAG-PKC signaling pathway by activating DGK activity (30). Remarkably, the treatment of cortical neuron cultures with PGZ at 10 μM for 1h normalized the phosphorylation of PKC in *Fmr1*-KO while its level in WT remained unaffected by the treatment (**Fig. 1B**). Several protein effectors of ERK pathway have been reported to be over-activated in neocortex of *Fmr1*-KO mouse (18) and in FXS patients (33). Consistent with an increased ERK activity, eukaryotic initiation factor 4E (eIF4E) phosphorylation is elevated in the brain of individuals with FXS and *Fmr1*-KO mice (26). Treatment of cortical neuron cultures with PGZ normalized phosphorylation of eIF4E in *Fmr1*-KO neurons while the WT level was non-significantly affected by the treatment (**Fig. 1C**).

**Figure 1:**
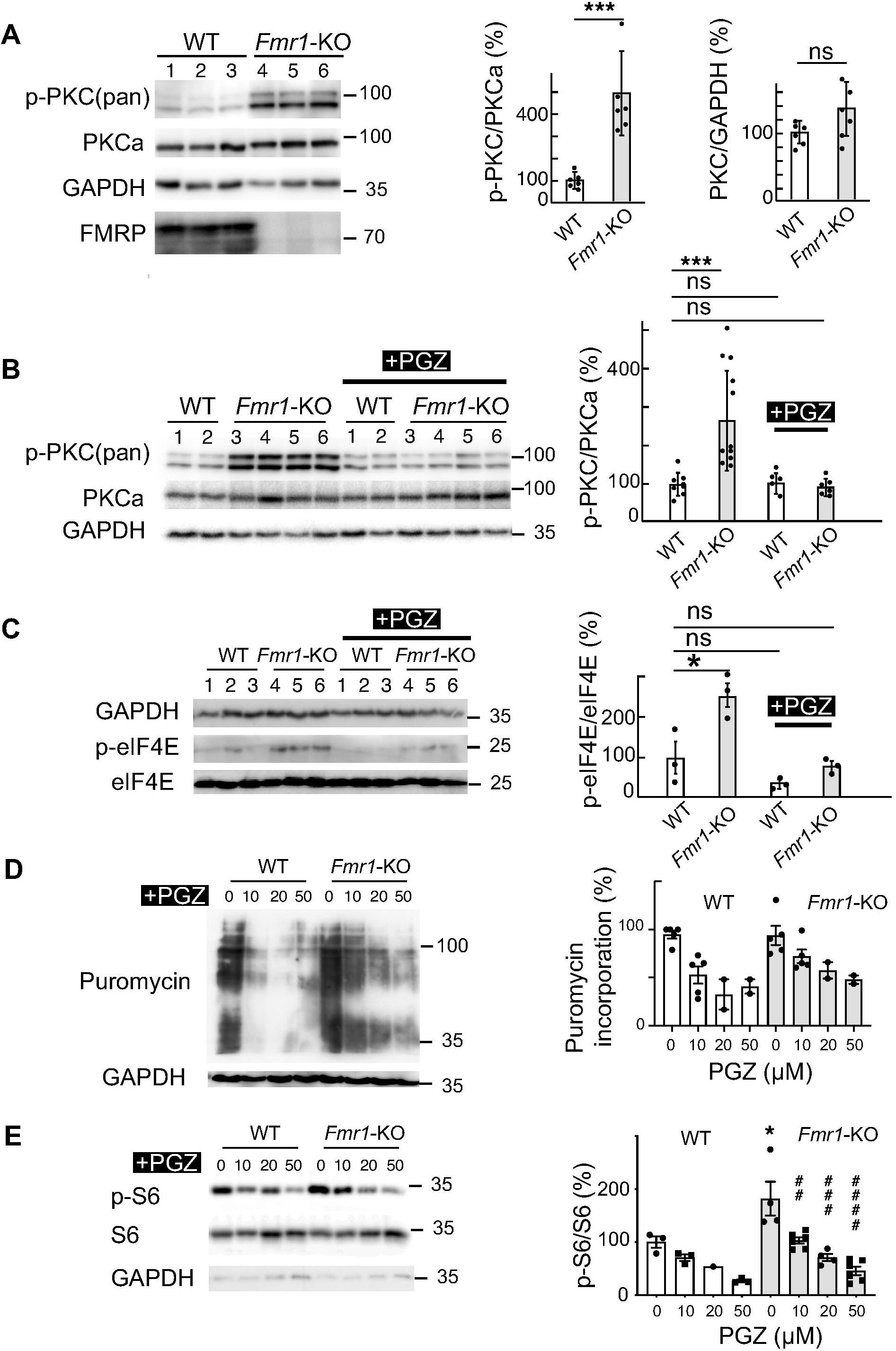
Pioglitazone corrects the excessive downstream DAG signaling in *Fmr1*-KO neurons. **A)** Phosphorylation of PKC is enhanced in *Fmr1*-KO compared to WT neurons. Representative immunoblots of lysates from WT and *Fmr1*-KO cortical neurons with the indicated antibodies (3 biological replicates are presented, molecular weight marker positions (kDa) are shown) and quantification of phosphorylation of PKC and total levels of PKCa. GAPDH was used as a loading control. For quantification, the phospho-protein signal was normalized first against PKCa protein signal and is presented relative to the signal of WT neurons (n = 6 embryos in each group). **B)** PGZ (10 μM, 1h) normalizes PKC phosphorylation of *Fmr1*-KO neurons. Representative immunoblots of lysates from WT and *Fmr1*-KO cortical neurons (2 biological replicates for WT and 4 for *Fmr1*-KO are shown). Quantifications were made as in A with n ≥ 6 embryos in each group. **C)** PGZ (10 μM, 1h) normalizes eIF4E phosphorylation of *Fmr1*-KO neurons. Representative immunoblots of lysates from WT and *Fmr1*-KO cortical neurons (3 biological replicates for WT and *Fmr1*-KO are shown). Quantifications of p-eIF4E signal were normalized first against GAPDH protein signal and are presented relative to the signal of WT with n = ≥ 5 embryos in each group. **D)** PGZ reduces protein synthesis rate in WT and *Fmr1*-KO neurons. Representative immunoblots of puromycin-labelled neuron lysates from WT and *Fmr1*-KO cortical neurons treated with pioglitazone for 1h at indicated doses. Quantifications of puromycin signal were normalized first against GAPDH protein signal and are presented relative to the signal of WT with n = 2 embryos. **E)** PGZ (24h) normalizes S6 phosphorylation of *Fmr1*-KO neurons. Representative immunoblots of lysates from WT and *Fmr1*-KO cortical neurons. Quantifications of p-S6 signal were normalized as in C with n = ? 3 embryos in each group. In A-E, each point represents data from an individual neuron culture, all values shown as mean ± s.e.m. ***p< 0.001, **p< 0.01, *p< 0.05 vs Wt, ## p< 0.01, ### p< 0.001, #### p< 0.0001 vs *Fmr1*-KO untreated; ns, not significant; in A calculated by Student test and in B-E by one-way ANOVA with Tukey’s multiple comparison test.

As a possible consequence of increased DAG signaling and eIF4E activation, *Fmr1*-KO mice exhibit elevated basal levels of protein synthesis rate in brain (34). The 10-20% increase of protein synthesis rate previously reported in hippocampal slices (27, 28) could not be confirmed in cortical dissociated neurons cultures, possibly because the variability between neuron cultures was in the same range of effect. We did however observe a reduction of the translation rate in both *Fmr1*-KO and WT neurons after PGZ treatment (**Fig. 1D**), with a more pronounced dose reduction in WT compared to *Fmr1*-KO, possibly linked to a higher basal translation rate in *Fmr1*-KO, confirming the ability of PGZ to reduce protein synthesis rate.

Not only ERK but also mTOR signaling pathways is hyper-activated in *Fmr1*-KO mice (26, 34, 35). Consistent with mTOR signaling activation, protein S6 phosphorylation is increased in *Fmr1*-KO neurons and this increase is normalized with PGZ treatment (**Fig. 1E**).

### Pioglitazone is delivered efficiently in brain of *Fmr1*-KO mouse by intraperitoneal injection

PGZ has been demonstrated to cross blood brain barrier (36) and, based on our above data, appeared as an interesting drug to test on *Fmr1*-KO mouse FXS model. To define the best route to deliver PGZ efficiently in brain, we compared several modes of administration: subcutaneous diffusing micropellet, food (solid pellets), daily intraperitoneal injection (IP). We chose a 20 mg/kg mouse/day dose as an average upper high dose in mouse based on literature survey. Treatment was started on 5-week-old mice and continued for three weeks. PGZ quantification in brain by HPLC-MS/MS revealed that IP administration enables to reach the highest dose of PGZ in the brain (19.4±9 nM) compared to food (3.7±0.9 nM) and subcutaneous administration (0.42±0.26nM), with no significant variation between *Fmr1*-KO and WT (**Fig. 2**). Based on these data we chose to administer PGZ via IP route.

**Figure 2:**
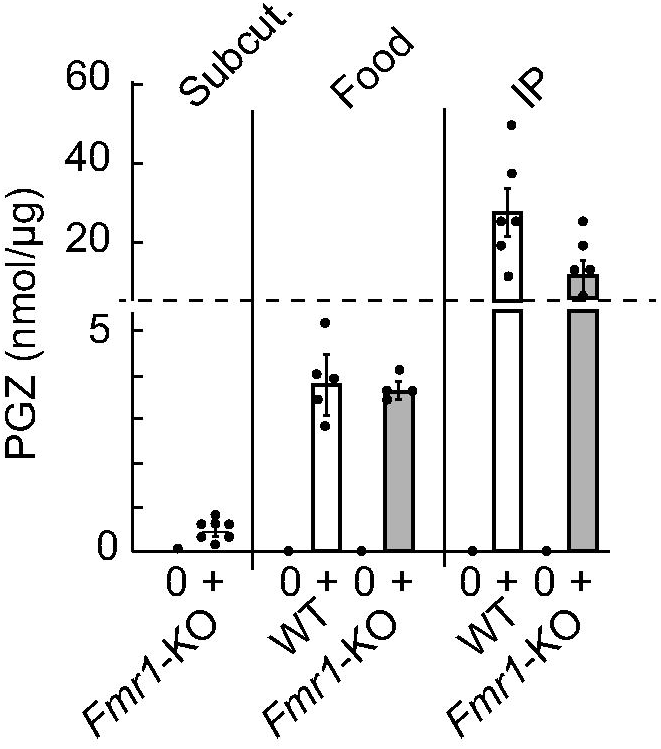
Pioglitazone concentration in total brain extracts. PGZ concentration (nM) was measured by mass spectrometry after subcutaneous (Subcut.), alimentation (food), or intraperitoneal (IP) administration (+) or mock treatment (0) in WT compared to *Fmr1*-KO n≥5.

### Pioglitazone daily IP treatment ameliorates *Fmr1*-KO behaviors, macro-orchidism and DAG

Five-week-old *Fmr1*-KO C57BL/6J hemizigous males and their wild-type (WT) littermates were injected IP with PGZ (20 mg per kg bodyweight per day) or vehicle for 7 days before starting the behavioral tests (**Fig. S1**) and IP injections were continued daily after each behavioral session (i.e. 19 d total). In the open field test, vehicle-treated *Fmr1*-KO mice did not show specific alterations compared to vehicle-treated WT mice in all the parameters analyzed (distance travelled, % time in center in arena, distance travelled in center of arena, number of rearings (**Fig. S2A-C**), suggesting that, in the conditions where the animals were handled, the influence of the genotype was not visible on the locomotor activity and anxiety of the animals. Noticeably, both WT and *Fmr1*-KO PGZ-treated mice had an average reduction of distance travelled and rearing numbers by approximately 20%, appearing as an unexpected previously unreported effect of the drug on mouse locomotor and rearing activity.

In the novel object recognition test, vehicle-treated *Fmr1*-KO mice showed a clear alteration of their ability to discriminate a novel vs a familiar object (recognition index = 49.89±1.33, not different from chance = 50%, **Fig. 3A**) compared to vehicle-treated WT (recognition index = 66.95±2.38). This difference was not the consequence of an altered exploration time of objects since this parameter was similar in both genotype and treatment groups (**Fig. S3A**). PGZ-treated *Fmr1*-KO mice had a fully rescued ability to discriminate the novel object (recognition index =68.33±2.28 not significantly different from WT 66.95±2.38). Treatment had no visible effect on the WT mice discrimination and did not modify the exploration parameters of both genotypes.

**Figure 3:**
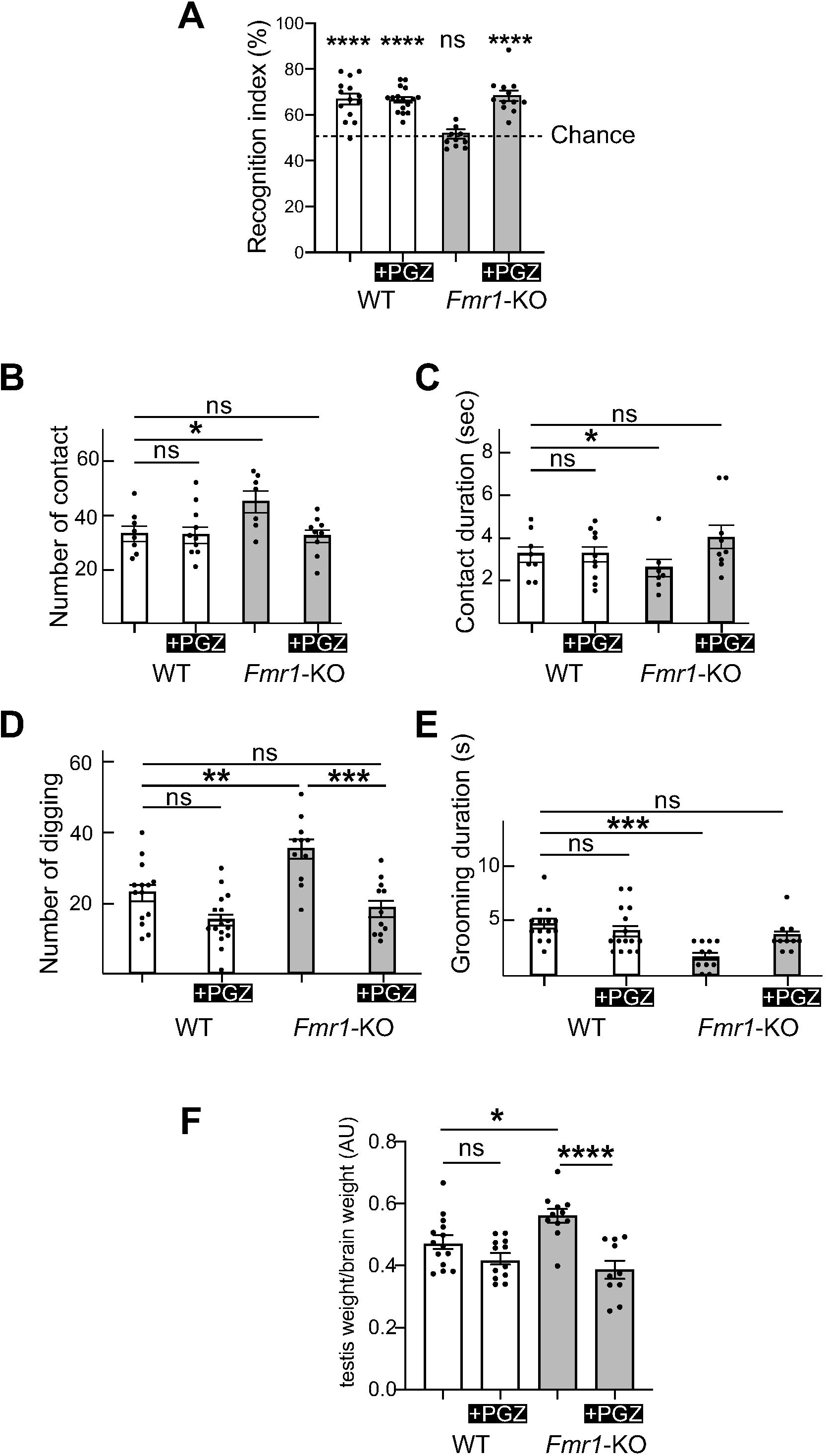
Pioglitazone corrects main phenotypes of *Fmr1*-KO mouse. **A**) Novel object recognition. Recognition index and duration of objects exploration during the acquisition and retention trials. Data are expressed as mean ± SEM and analyzed using one-sample t, **** p<0.0001 vs chance (50%), one outlier in *Fmr1*-KO vehicle was removed (ROUT Q=1%). **B, C**) Free sociability dyadic reciprocal social interaction test. Number (**B**) and duration (**C**) of contacts. Data are expressed as mean ± SEM and analyzed using one-way ANOVA and Tukey’s multiple comparisons test. * p<0.05 vs WT vehicle. Number of digging (**D**) and digging duration (**E**). Data are expressed as mean ± SEM and analyzed using one-way ANOVA and Tukey’s multiple comparisons test. **** p<0.0001, *** p<0.001, ** p<0.01, ns non-significant. **F**) Pioglitazone corrects macro-orchidism. Mean testicular weight of vehicle- and PGZ-treated WT and *Fmr1*-KO mice. Data are expressed as mean ± SEM and analyzed using one-way ANOVA and Tukey’s multiple comparisons test, **** p<0.0001, * p<0.05, ns non-significant.

In the free sociability dyadic reciprocal social interaction test, vehicle-treated *Fmr1*-KO mice showed an increased number of contacts with shorter contact duration time compared to WT mice (**Fig. 3B, C**). In contrast, PGZ-treated *Fmr1*-KO mice were not different from vehicle-treated WT, suggesting that treatment had normalized their social behavior. In contrast, vehicle-treated WT mice were not visibly behaving differently from their PGZ-treated WT littermates in this test.

Vehicle-treated *Fmr1*-KO mice showed a significant increase of number of digging compared to vehicle-treated WT (**Fig. 3D**) while PGZ-treated *Fmr1*-KO mice were not different from vehicle-treated WT suggesting also that treatment had normalized their digging behavior. PGZ-treated WT showed a trend to have a decreased number of digging compared to vehicle-treated WT but this was not significant. Vehicle-treated *Fmr1*-KO mice showed a significant decrease of grooming duration compared to vehicle-treated WT (**Fig. 3E**) while PGZ-treated *Fmr1*-KO mice were not different from vehicle-treated WT suggesting also that treatment had normalized their behavior. PGZ-treated WT showed no difference compared to vehicle-treated WT.

In the rotarod test, no influence of the genotype or of the treatment was observed, neither in habituation nor in the following sessions (**Fig. S3B**). Prepulse inhibition test was performed but there was no significant difference between genotypes and the treatment groups (**Fig. S3C**), but this test showed variability across the studies (37-39), possibly related to the sensitivity of recording methods used.

Macro-orchidism is a hallmark of post-adolescent male individuals with FXS (1) and is present in *Fmr1*-KO mice (40). PGZ administration for 19 d reduced the testicular weight in *Fmr1*-KO mice to the same level as the WT treated mice (**Fig. 3F**). Noticeably, PGZ had a trend to reduce the testicular weight in WT mice, suggesting a possible impact of PGZ both in normal and pathological context. Meanwhile, no significant difference for total or brain weight was observed between genotype and treatment groups (**Fig. S4A,B**). PGZ treated animals showed a trend to have a reduced weight gain during the time of tests, but this was not statistically significant. Noticeably, while the body composition of vehicle-treated *Fmr1*-KO mice was not significantly different from vehicle-treated WT mice, PGZ treatment induced a significant increase of water retention in the tissues of *Fmr1*-KO mice (**Fig. S4C-G**), a well-known side effect of PGZ treatment (41). Diacylglycerol (DAG) is increased in *Fmr1*-KO cortical neurons and in cerebellum of FXS patient, in agreement with a loss of DGKk activity in FXS neurons (8). The level of several main DAG species was found to be increased in vehicle-treated *Fmr1*-KO total brain extracts, notably main DAG species 38:4, and their level was normalized in PGZ-treated *Fmr1*-KO mice (**Fig. 4**).

**Figure 4:**
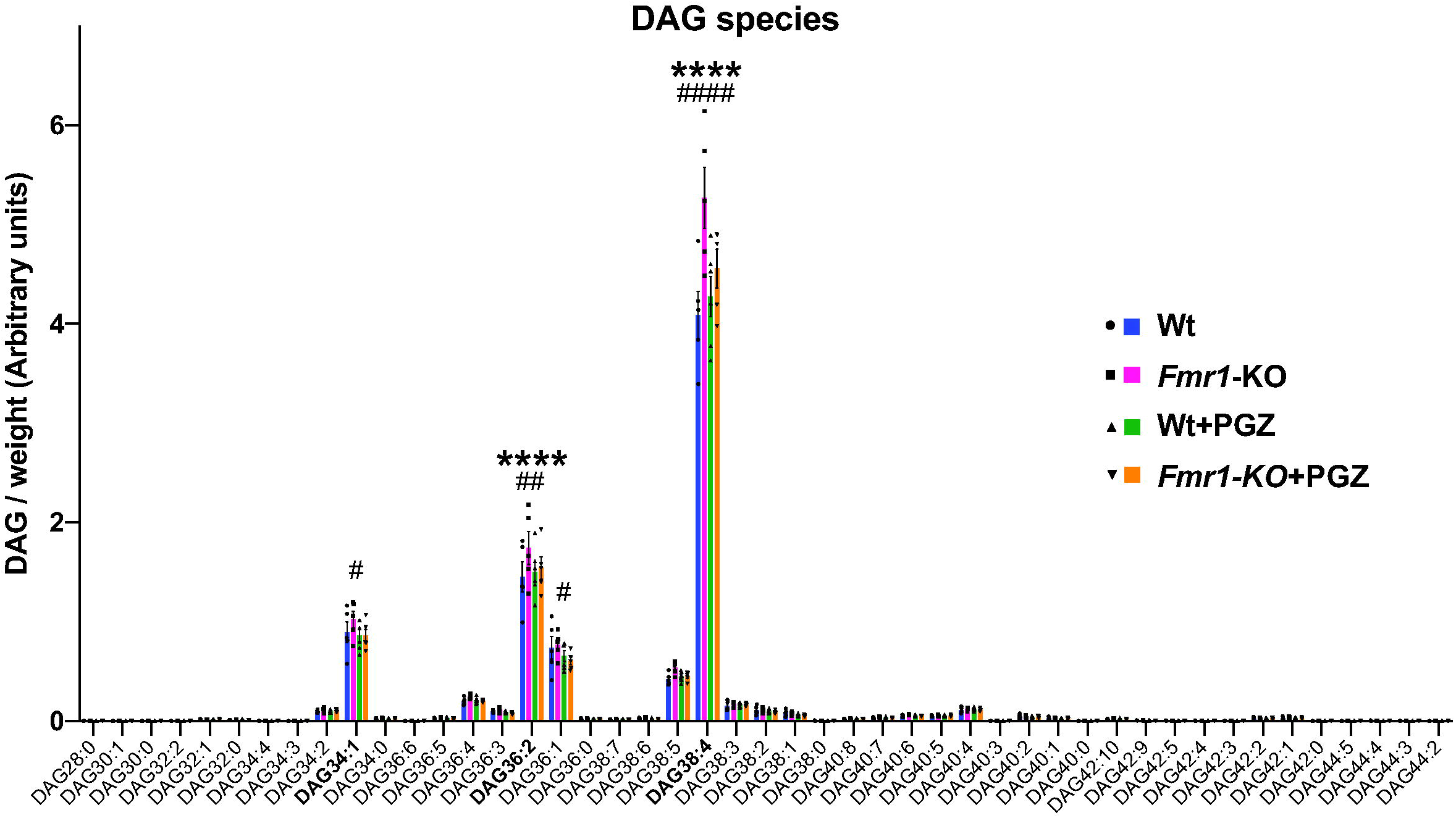
Pioglitazone corrects elevated DAG level in brain of *Fmr1*-KO mouse. Measure of diacylglycerol species in WT and *Fmr1*-KO mouse total brain extracts by LC-MS/MS. The level of DAG species analyzed by LC-MS/MS is expressed as arbitrary units relative to DAG standard added to samples prior lipid extraction and normalized to weight of samples. An increase of most abundant DAG species (36:2 and 38:4) is observed in *Fmr1*-KO compared with WT neurons. Data are expressed as mean ± SEM and analyzed using ordinary 2way ANOVA and Tukey’s multiple comparisons test, n = 5-6 biological replicates, ****P < 0.0001 *Fmr1*-KO vs WT, ####P < 0.0001, ##P < 0.01, #P < 0.05 *Fmr1*-KO +Pio vs *Fmr1*-KO.

## Discussion

In this study, we tested the ability of diacylglycerol kinase agonist pioglitazone (PGZ) to correct the symptoms of *Fmr1*-KO mouse model of fragile X syndrome. The rationale of this study is based on the fact that diacylglycerol kinase kappa (DGKk), a master regulator whose mRNA is a main target of FMRP in neurons, is under-expressed in the brain of *Fmr1*-KO mouse and signs of reduction of DGK activity (namely DAG/PA imbalance) are observed both in *Fmr1*-KO mouse and in FXS patients (8). Substrate of DGKk, DAG, is a prominent modulator of synaptic transmission probably in all types of synapses (42-44) that is produced by phospholipase C (PLC) from abundant phosphatidylinositol 4,5-bisphosphate (PIP2) membrane stores (45) after stimulation of G protein-coupled receptors such as mGluRI. The excess of DAG originating from the loss of DGKk activity could play a prominent role in FXS pathomechanism because DAG, through activation of its main effector protein kinase C (PKC) (32), could be responsible of mitogen activated protein kinase (MAPK/ERK) (46) or Ras/phosphatidylinositol-3-kinase (PI3K)/AKT pathways (47). These pathways have been demonstrated to be overactivated in *Fmr1*-KO mice (34, 35, 48, 49). In agreement with a neuronal DAG increase in FXS condition, we showed that PKC is overactivated in *Fmr1*-KO neurons (**Fig. 1A**). Overall, the excess of DAG resulting from the lack of DGKk activity in *Fmr1*-KO mouse could underlie the molecular mechanism leading to the excessive mGluRI signaling and the increase of local protein synthesis (29, 50). Based on the hypothesis that DGK activity alteration could contribute to the severity of FXS condition, we postulated that increasing DGK activity could be beneficial for the patients. We tested the impact of DGK agonist pioglitazone (PGZ) on *Fmr1*-KO mouse model and showed that PGZ corrects its molecular and behavioral alterations. In cortical neuron cultures, PGZ normalized the overactivation of PKC and eIF4E (respectively first and downstream effectors of DAG signal transduction pathway) observed in *Fmr1*-KO compared to WT neurons (**Fig. 1B,C**). These data suggest that PGZ is able to inhibits DAG-PKC signaling in neurons like in endothelial cells (30). PGZ demonstrated also the ability to reduce the protein synthesis rate in neurons cultures (**Fig. 1D**). Thus, PGZ appears to be able to counteract some of the well-established molecular alterations of *Fmr1*-KO neurons. The administration of PGZ via intraperitoneal injection resulted in a better concentration of the molecule in the brain compared to an administration through alimentation (3.7±0.9 vs 19.4±9 nM, **Fig. 2**), however we cannot exclude that a proportion of the molecule was degraded during the food pellet preparation process that involved heating at 60°C for several minutes. Subcutaneous pellets that enable slow diffusion of the molecule while avoiding repeated injections appeared to be inappropriate for targeting the brain, giving the lowest concentration in brain.

PGZ administered daily for 7 days prior the beginning of behavioral tests and continued during the tests for a total of 19 days at 20 mg/kg/day (corresponding to approximately half the maximum recommended human oral dose of 45 mg based on mg/m^2^, Takeda Inc.) was able to correct the elevated level of main DAG species in the brain of *Fmr1*-KO mouse (**Fig. 4**) in agreement with its DGK agonist effect (30).

PGZ normalized the observed behavioral alterations of the *Fmr1*-KO model. Notably, the recognition of novel object after 24h that was absent in *Fmr1*-KO was reestablished in *Fmr1*-KO-treated mice (**Fig. 3A**), suggesting that PGZ had effect on long term memory deficit of the FXS model. Reciprocal social interaction, stereotypical behavior (digging) and grooming were all normalized (**Fig. 3B-D**). We did not observe influence of genotype on locomotor activity but unexpectedly PGZ decreased the distance traveled and rearing numbers in both genotype, suggesting a possible adverse effect of treatment on voluntary locomotor activity (**Fig. S2A**). Whether linked or not with this effect, PGZ had an influence on free body fluid retention, this is a well-known edema causing effect of the molecule due to renal sodium retention and vascular hyperpermeability (51). However, the physical performances as assessed with rotarod test were not affected.

PGZ inhibits DAG-PKC signaling by activating DGK enzymes (30), but the mechanism of this activation is still unclear. PGZ can act by PPAR gamma dependent and independent pathways (52). In PGZ-treated neuron cultures and in brain extracts of PGZ-treated mice, no change was observed on the transcript level of DGK enzymes (**Fig. S5**) suggesting that in neurons, unlike in endothelial cells (30), PGZ mediates DGK activity in a PPAR gamma independent manner. This is in agreement with the fact that PPAR gamma is highly expressed in adipose tissues (53) but very poorly expressed in brain (54). The exact mechanism of PGZ action in the brain remains thus to be understood.

An augmentation of cerebral DAG has been observed to coincide with the appearance of mild cognitive impairment in Alzheimer’s (AD) (55), Parkinson’s and Lewy Body disease (56) and this increase has been proposed as a triggering factor of neuroinflammation (57). PGZ has established anti-inflammatory effects and had been shown to have positive cognitive effects on mouse models of AD (58, 59), that like the FXS model have aberrantly elevated glutamatergic neurotransmission (60). Neuroinflammation has been evoked to play a role in FXS and other form of ID and autism (61). In fact, GSK3, a strong promoter of inflammation in various tissues (62) has been shown to be overactivated in *Fmr1*-KO brain (63). GSK3 activity has been demonstrated to be trigered by DAG/PA balance, notably by its impact on PI4P5-kinase to activate AKT/phosphatidynositol 3-kinase (PI3K) controlling GSK3 activity status (64, 65). Moreover, knock-out of DGKß, an isozyme of DGKk also expressed in brain, leads to an increase of GSK3ß activity proposed to contribute to the hyperactivity and reduced anxiety observed in the model (66). Thus, the loss of DGKk activity could be also responsible for this type of neuroinflammation related phenotypes and PGZ may also contribute to interfere with these alterations.

In conclusion, in this study we provide lines of evidence that pioglitazone, one of the most widely used US Food & Drug Administration (FDA)-approved anti-diabetic corrects main molecular and phenotypic deficits of the FXS *Fmr1*-KO model. To our knowledge, this is the first study to report that acting on the DAG signaling in the brain can correct the behavioral alterations associated with FXS. Upregulating DGK activity may thus represent a potential therapeutic strategy for the treatment of FXS.

## Acknowledgements

We thank Dr. F. Sakane, Dr. R. Willemsen, Dr N. Vitale, Dr F. Riet for materials and discussions, N. Banquart-Ott and C. Nahy for mice care. PGZ analyses were performed at the Plant Imaging & Mass Spectrometry, PIMS, CNRS Strasbourg. We are very grateful to all colleagues of the team for discussions and suggestions. The DAG analyses were performed at the Metabolome Facility-MetaboHUB (ANR-11-INBS-0010). This work was supported by ANR-18-CE12-0002-01 and Fondation Jérôme Lejeune funding to HM, by FRAXA Foundation post-doc fellow to AG and by Fond Paul Mandel to KH. This study was also supported by ANR-10-LABX-0030-INRT under the frame program Investissements d’Avenir ANR-10-IDEX-0002-02.

## Author contributions

AG and KH performed most experiments and analyzed the data. DH LS and JZ performed PGZ mass spec analyses. LF performed diacylglycerol analyses. BZ, EF, assisted with molecular biology work. AD and YH helped with behavioral analyses. HM supervised the study and wrote the manuscript with AG, KH, BZ. All authors read the manuscript and approved the content.

**Correspondence and requests for materials** should be addressed to HM.

## Supplementary materials

**Table S1.**
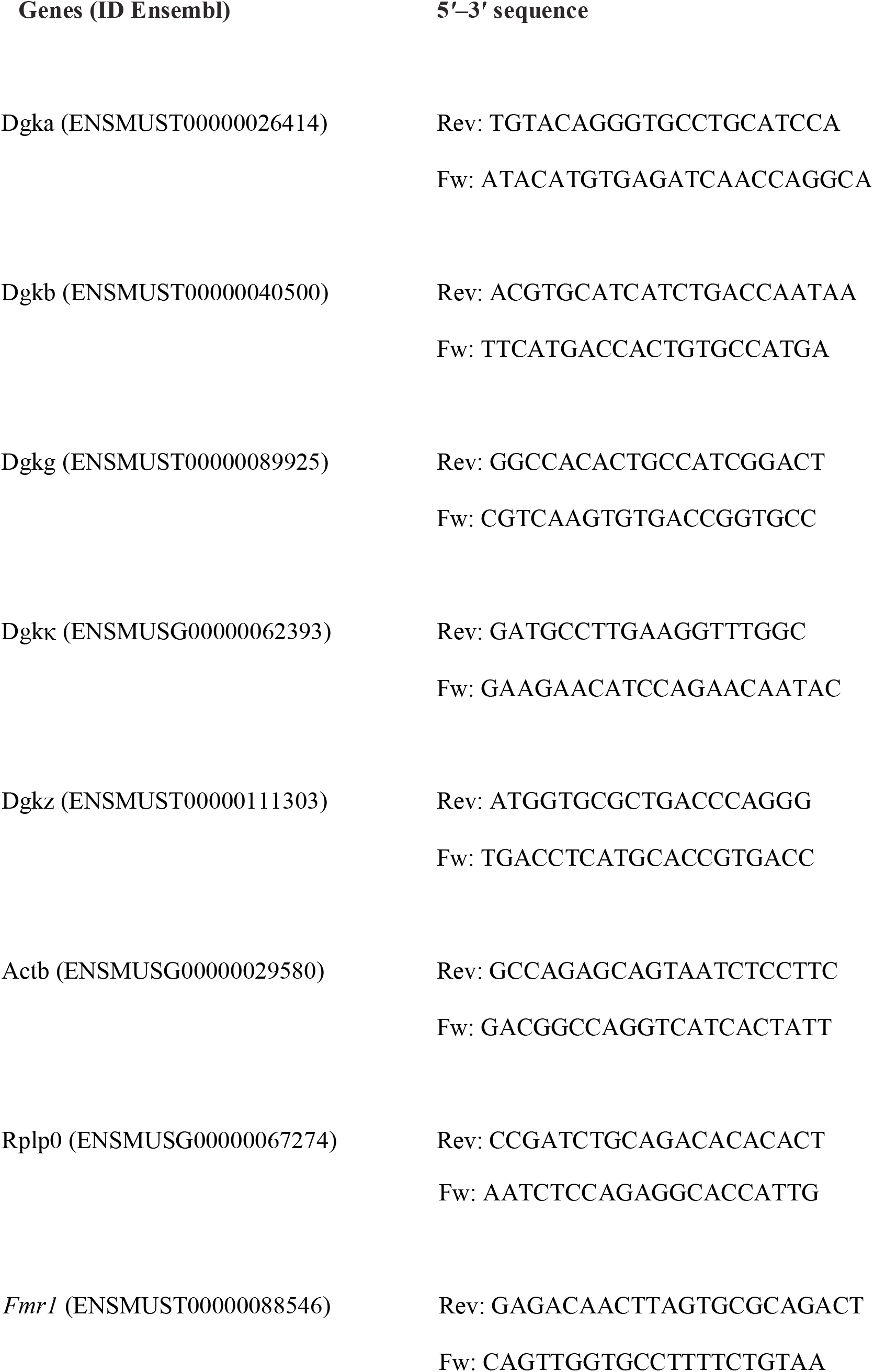
Primers used for qRT-PCR analysis.

**Figure S1: Phenotyping scheme**

**Figure S2: Influence of pioglitazone on locomotor activity in open field test. A)** Locomotor activity (distance travelled) in the whole arena, **B)** number of rearings and **C)** percentage of time spent in the center of the bowl during 30 min. Data are expressed as mean ± SEM and analyzed using one-way ANOVA and Tukey’s multiple comparisons test. * p<0.01 vs WT vehicle, ## p<0.01, # p<0.05 vs *Fmr1*-KO vehicle; ns, non-significant.

**Figure S3: Influence of pioglitazone on novel object recognition, rotarod, prepulse inhibition tests. A**) Novel object recognition. Mean sniffing duration per object passage (sec) during acquisition and retention phases. **B**) Rotarod test. Latency to fall during 4 training sessions. **C**) Acoustic startle reflex (ASR) and prepulse inhibition (PPI) analysis (BN, background noise; P*x*, stimuli of *x*dB such as P70 = 70 dB; ST110, acoustic startle pulse of 110 dB; %PP*x*, prepulse of *x*dB followed by a pulse of 110 dB, Glb, global fit). A, B, C, data are expressed as mean ± SEM and analyzed using one-way ANOVA and Tukey’s multiple comparisons test.

**Figure S4: Influence of pioglitazone on weight and body composition. A)** Weight evolution of vehicle- and PGZ-treated WT and *Fmr1*-KO mice throughout treatment (D, day of treatment). **B)** Mean brain weight of vehicle- and PGZ-treated WT and *Fmr1*-KO mice. **C)** Body composition determined by quantitative nuclear magnetic resonance (qNMR). Lean, free body fluid (FBF), fat and rest (other) are represented as % of the total body weight (means of n=3) and as total weight (g) **D, E, F, G**, respectively, expressed as mean ± SEM and analyzed with one-way ANOVA and Tukey’s multiple comparisons test, ** p<0.01 vs *Fmr1*-KO vehicle.

**Figure S5: Influence of pioglitazone on DGK transcripts level in total brain extracts and cortical neuron cultures.** The level of indicated DGK mRNAs was determined by qRT-PCR on total mRNAs extracted from **A)** total brain homogenates of mice treated with vehicle or PGZ 20mg/kg/day during 21 days and **B)** from 15 DIV cortical neuron cultures treated with 20μM PGZ for 1h. Fold change of expression was calculated by using the ΔΔCT method and with gene ActB as normalizer. Error bar is SD, n = 6 biological replicates.

