## Supplementary figures and images for "Pioglitazone improves deficits of *Fmr1*-KO mouse model of Fragile X syndrome by interfering with excessive diacylglycerol signaling"

### Supplemental Figure 1

**Fig. S1**

Geoffroy, Habbas et al.

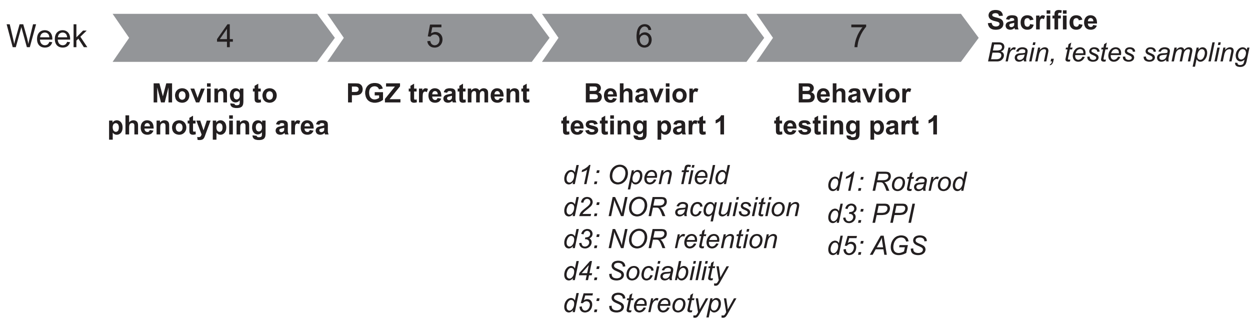

### Supplemental Figure 2

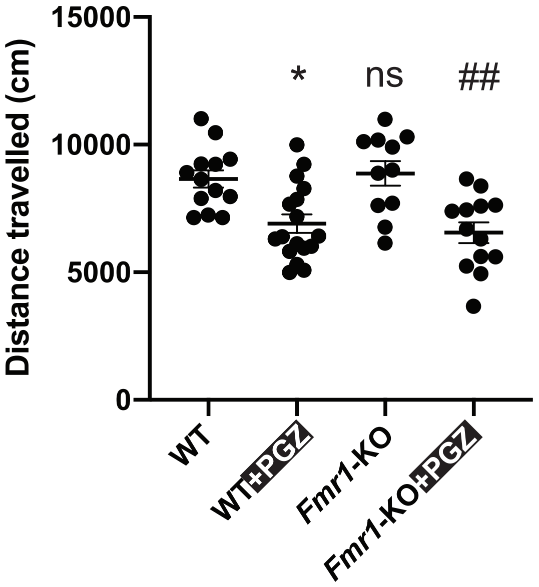

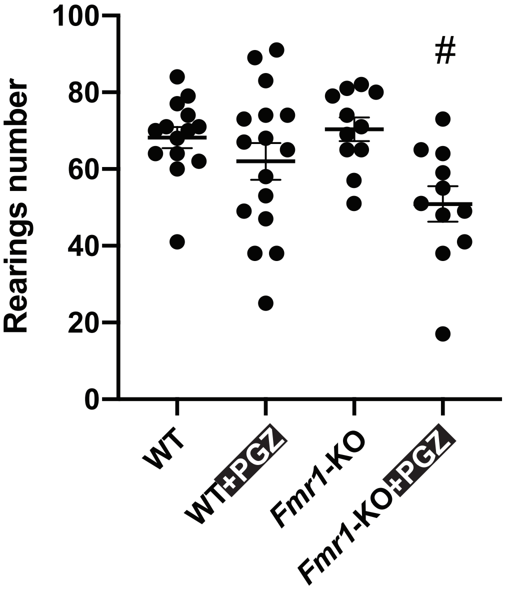

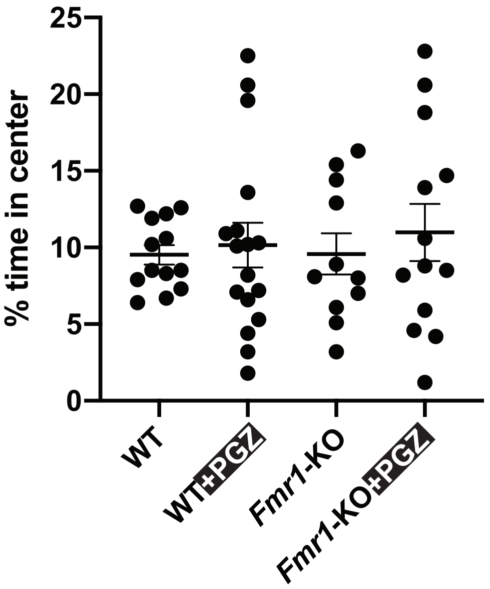

### Supplemental Figure 3

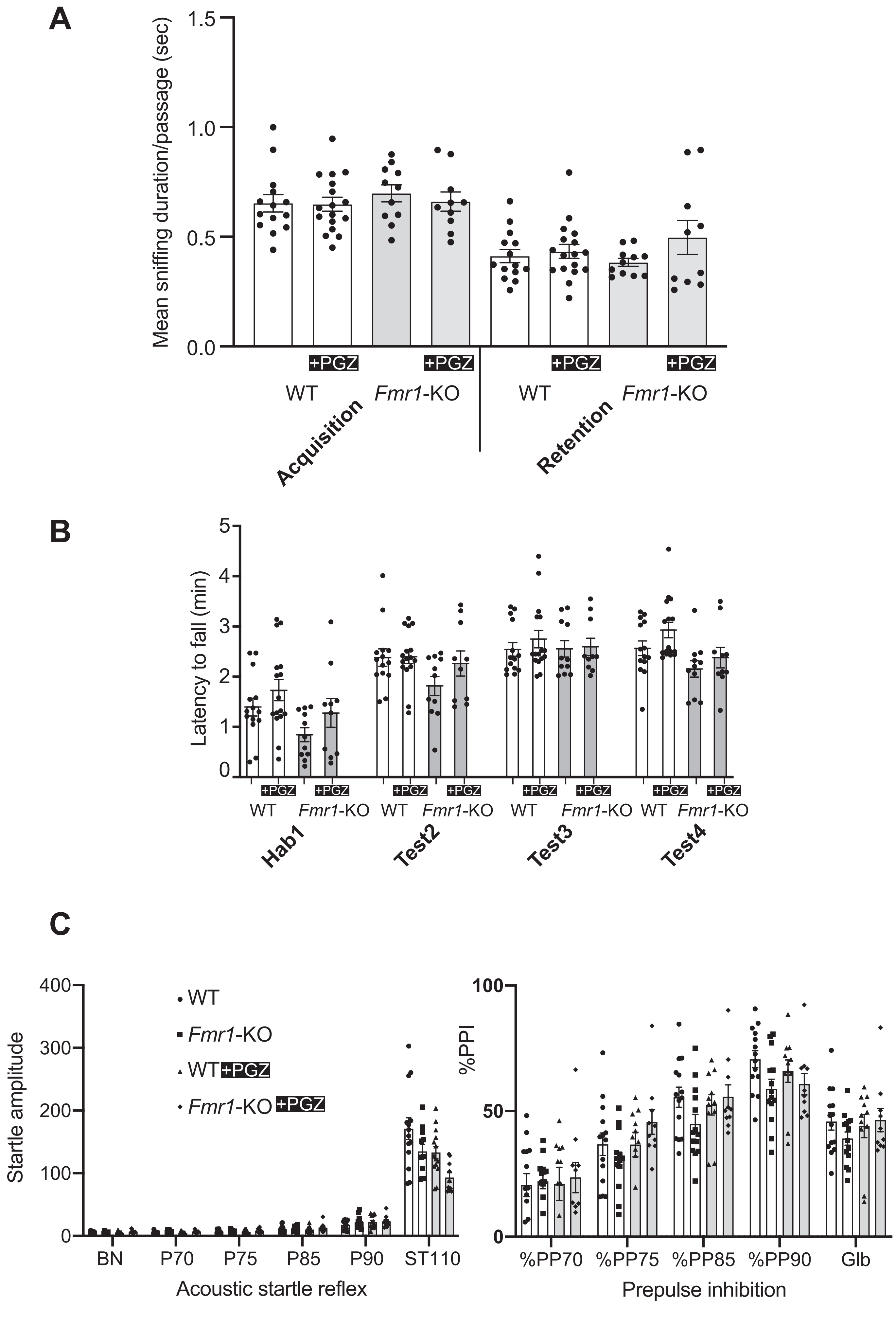

### Supplemental Figure 4

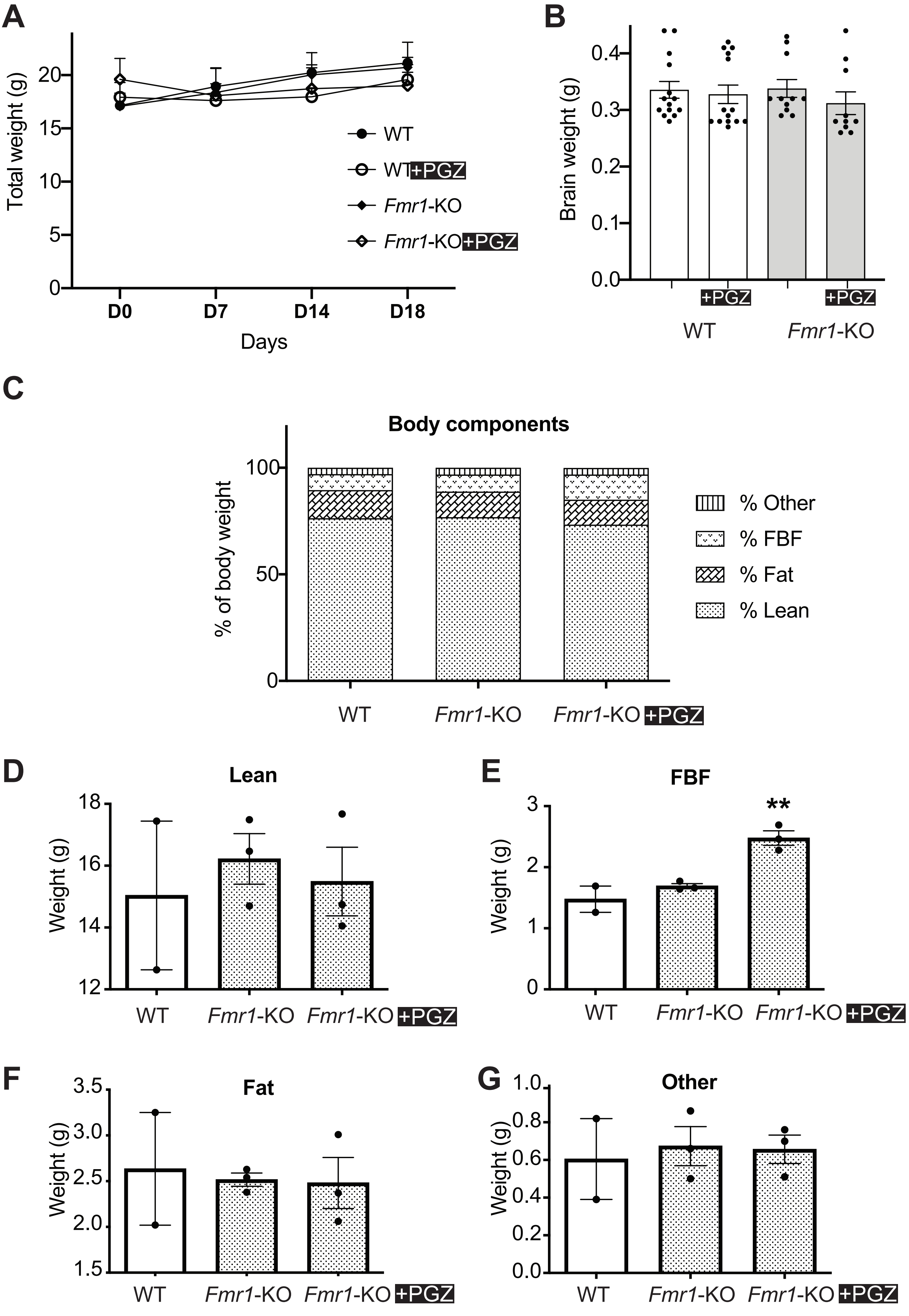

### Supplemental Figure 5

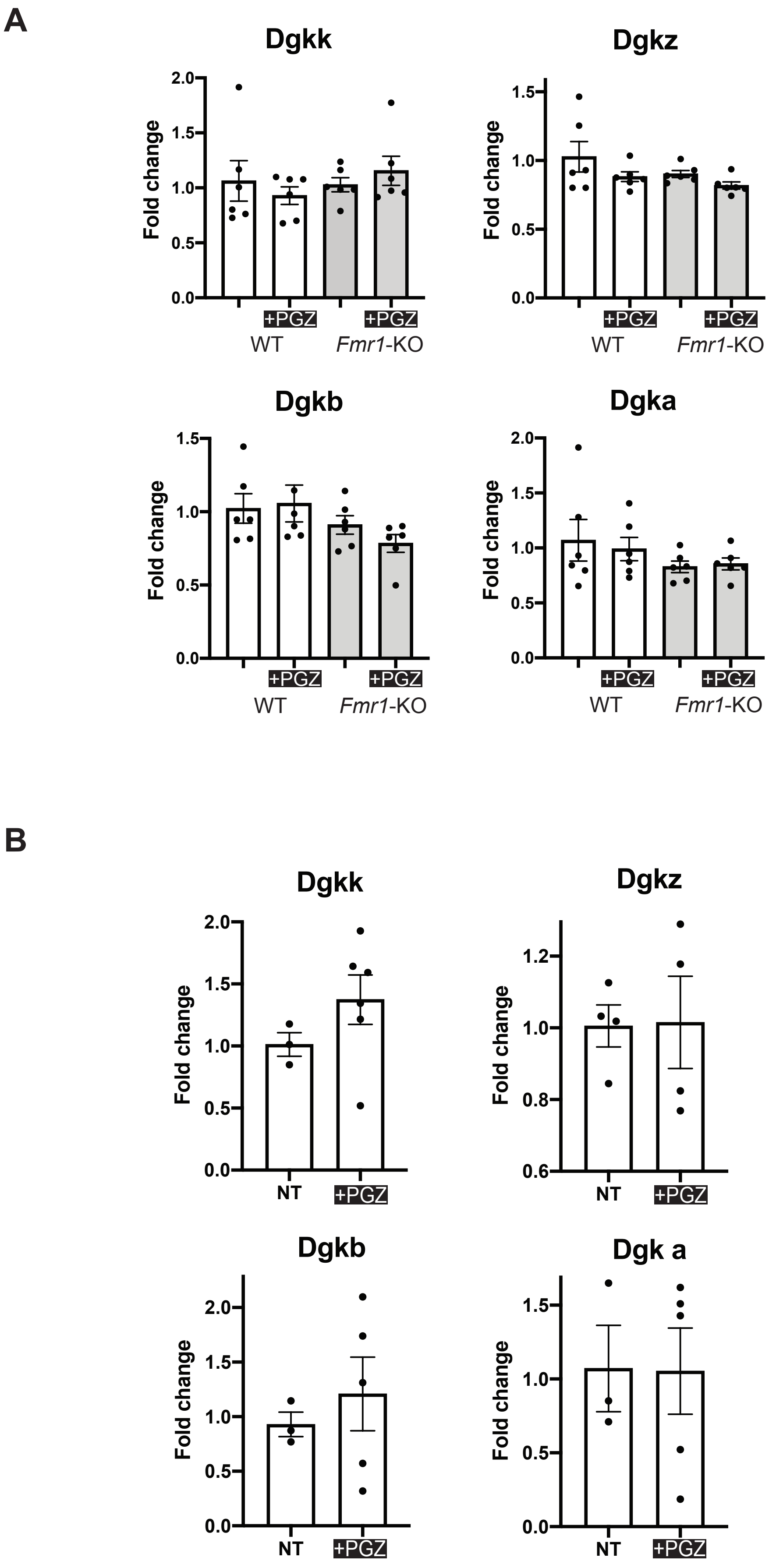
